# Arabidopsis root responses to salinity depend on pectin modification and cell wall sensing

**DOI:** 10.1101/2020.12.18.423458

**Authors:** Nora Gigli-Bisceglia, Eva van Zelm, Wenying Huo, Jasper Lamers, Christa Testerink

**Affiliations:** Laboratory of Plant Physiology, Wageningen University and Research, Wageningen 6708 PB, The Netherlands

**Keywords:** Salt stress responses, Plant cell wall signaling, pectin modifications, Cell wall integrity, *Catharanthus roseus* Receptor like Kinase 1 Like, THESEUS1, HERKULES1, FERONIA

## Abstract

Soil salinity is an increasing worldwide problem for agriculture, affecting plant growth and yield. To understand the molecular mechanisms activated in response to salt in plants, we investigated the *Catharanthus roseus* Receptor like Kinase 1 Like (CrRLK1L) family, which contains well described sensors previously shown to be involved in maintaining and sensing the structural integrity of the cell walls. We found that *herk1the1-4* double mutants, lacking the function of the *Arabidopsis thaliana* Receptor like Kinase HERKULES1 combined with a gain-of-function allele of THESEUS1, phenocopied the phenotypes previously reported in plants lacking FERONIA (FER) function. We report that both *fer-4* and *herk1the1-4* mutants respond strongly to salt application, resulting in a more intense activation of early and late stress responses. We also show that salt triggers de-methyl esterification of loosely bound pectins, responsible for the activation of several salt response signaling pathways. Addition of calcium chloride or chemically inhibiting pectin methyl esterase (PME) activity reduced activation of the early signaling protein Mitogen Activated Protein Kinase 6 (MPK6) as well as amplitude of salt-induced marker gene induction. MPK6 is required for the full induction of the salt-induced gene expression markers we tested. The sodium specific root halotropism response on the other hand, appears independent of MPK6 or calcium application, and is only mildly influenced by the cell wall sensors FER/HERK1/THE1-4 or alteration of PME activity. We hypothesize a model where salt-triggered modification of pectin requires the functionality of FER alone or the HERK1/THE1 combination to attenuate salt responses. Collectively, our results show the complexity of salt stress responses and salt sensing mechanisms and their connection to cell wall modifications, responsible for several salt response pathways and ultimately plant resilience to salinity.

## Introduction

For decades, researchers have tried to discover how plants sense salt. Salinity of the soil is highly detrimental to plant growth as salt simultaneously causes both ionic and osmotic stress, triggering subsequential general and specific signaling responses (van Zelm et al., 2020). Recently, the role of plant cell wall modifications as possible factors in regulating salt stress responses have gained special interest (Feng et al., 2018). In plants, cell walls act as a barrier that protects plants to environmental threats and functions as structural support for the whole plant body (Underwood, 2012). In young Arabidopsis plants, primary cell walls (CWs) are made of complex polysaccharide networks where the main load bearing component is cellulose, a linear polymer composed of amorphous and crystalized β 1-4 linked glucose monomers. Cellulose fibrils are intimately connected with pectin, another major component of the cell wall, described to be essential for regulating cell expansion and elasticity (reviewed in (Gigli-Bisceglia et al., 2020)). Contrary to cellulose, which is synthetized at the plasma membrane level, pectins are synthetized in Golgi bodies and released in the apoplast in a highly methyl-esterified form (Ridley et al., 2001; Willats et al., 2001). Cell walls are subjected to controlled modifications that allow the cell to extend, driving the direction of cell growth (reviewed in (Vaahtera et al., 2019)). Cell wall modifications also happen in response to both biotic and abiotic stresses and these structural changes are perceived and sensed by plants. Receptors/sensors are responsible for triggering intracellular downstream signals in response to turgor pressure driven cell wall-dependent mechanical distortions, a process called cell wall integrity (CWI) maintenance (Levin, 2011; Rui and Dinneny, 2020).

Several members of the *Catharanthus roseus* Receptor Like Kinase1 Like (CrRLK1L) family have been described to play a role in mechanisms related to cell wall remodeling (*e.g.* cellulose impairment, pathogen infections, fertility) (Nissen et al., 2016). CrRLK1Ls contain 2 malectin domains, proposed to selectively bind complex carbohydrates, and an intracellular kinase domain responsible for transmitting signals intracellularly (Schallus et al., 2008). In Arabidopsis, 17 genes encoding for CrRLK1Ls have been identified and in some cases their functional roles have extensively been studied. For example FERONIA (FER), appears to be vital in multiple aspects of plant life (*e.g.* fertility, root hair development, mechanical stress, immune responses) while THESEUS1 (THE1), HERKULES 1/2 (HERK1/2) and CURVY (CVY1) are essential for cell elongation and cell morphology related processes (Guo et al., 2009; Nissen et al., 2016; Voxeur and Höfte, 2016).

While they are extensively studied, the mechanism through which CrRLK1Ls work is not fully understood. It seems that CrRLK1Ls might form complexes through transient hetero-oligomerization as found for the newly identified LETUM1/LETUM2 modulating plant autoimmunity and HERK1/ANJEA or ANXUR (ANX)/BUDDHA’S PAPER SEAL (BUPS) that control pollen tube related mechanisms (Ge et al., 2019; Huang et al., 2020). CrRLK1Ls also act as receptors for small extracellular peptides called Rapid Alkalinization Factors (RALFs). In Arabidopsis, several different *RALF* genes have been identified and RALF22, 23, 33, 34 and RALF1 have been reported to function as FER ligands, with the latter being required for Arabidopsis ovule fertilization (Abarca et al., 2020; Bedinger et al., 2010; Ge et al., 2019; Gonneau et al., 2018). On the other hand, THESEUS1 (THE1) seems to be the receptor of RALF34, and both (THE1/RALF34) are essential for regulating lateral root development in Arabidopsis (Gonneau et al., 2018). Some CrRLK1Ls have been defined being responsible for sensing cell wall alterations and mechanical stress responses with THESEUS1 (THE1) being essential to fully activate at least one branch of the CWI signaling pathways (Engelsdorf et al., 2018; Gigli-Bisceglia et al., 2018). In Arabidopsis several other proteins have been characterized to play a role in CWI signaling, including stretch-activated plasma membrane-localized calcium channels (MCAs) and the Nitrate Reductase 1 and 2 (NIA1/2), the latter being recently described to be play an essential role in the cell wall damage dependent cell cycle regulation (Engelsdorf et al., 2018; Gigli-Bisceglia et al., 2020, 2018; Vaahtera et al., 2019).

Interestingly, many of the CWI players have been found to overlap with salt stress response regulators. This is the case for Leucin Rich Repeat Receptor Like Kinases MIK2/KISS (MALE DISCOVERER 1-INTERACTING), FEI1/2, FERONIA, but also for NIAs and MCAs (Feng et al., 2018; Harpaz-Saad et al., 2012; Shabala et al., 2015; Van der Does et al., 2017; Xie et al., 2013). Intriguingly, while loss of function mutants for *mik2, fei1/2, nia1/2* and *mcas* (*mca1,2*) were reported to be highly sensitive to salt application, their responses to chemically-induced cellulose impairment varied, with *fei1/2* being extremely sensitive and *mik2, nia1/2* and *mca1,* being partly or almost completely insensitive to cellulose inhibition (Engelsdorf et al., 2018; Gigli-Bisceglia et al., 2018; Harpaz-Saad et al., 2012; Shabala et al., 2015; Van der Does et al., 2017; Xie et al., 2013). Yet how and why CWI and salt induced signaling overlap has been poorly investigated.

Within seconds to minutes after salt treatment, early signaling molecules are induced, such as calcium spikes, reactive oxygen species (ROS) accumulation and mitogen protein kinase (MAPK) cascade phosphorylation (reviewed in (Lamers et al., 2020; van Zelm et al., 2020)). Only recently, the identification of MONOCATION-INDUCED [Ca^2+^]_i_ INCREASES 1 (MOCA1) mutant phenotype has highlighted that modification of PM-localized sphingolipids can alter salt sensitivity. MOCA1 previously known as IPUT1, functioning as glucuronosyltransferase and essential to transfer glucuronic acid (GlcA) to inositol phosphorylceramide (IPC) sphingolipids (named GIPCs), has been described important for regulating GIPCs content (Jiang et al., 2019). GIPCs have been suggested to bind to monovalent cations (such as Na^2+^), triggering the opening of unknown plasma membrane localized Ca^2+^ channels involved in calcium spikes (Jiang et al., 2019). The accumulation of intracellular calcium is necessary to activate the salt-overly-sensitive (SOS) pathway, which consists of the Na^+^/H^+^ exchanger SOS1 and SOS2/3 signaling components, all required for limiting the damaging effects of salt on cells by increasing Na^+^ efflux (Ji et al., 2013; Yin et al., 2020; Zhou et al., 2016). On the other hand, the change in main root direction observed in high salt concentrations, called halotropism, was actually stronger in the *sos* mutants, and was shown to be dependent on auxin distribution (Deolu-Ajayi et al., 2019; Galvan-Ampudia et al., 2013). Interestingly, salt-stress induced calcium spikes depend on FER, as *fer-4* loss of function mutants are unable to induce calcium bursts as wild type plants (Feng et al., 2018). FERONIA seems to be required to maintain cell wall architecture in response to salt stress since loss of function *fer-4* mutants display cell bursting and severe cell swelling in the root in the presence of high salt concentrations.

It has been proposed that the salt oversensitive phenotypes observed in FER mutants depend on the enhanced cell wall softening due to the presence of cell wall localized sodium ions. These results are corroborated by supporting data showing that FER’s malectin domain can bind pectin macromolecules *in vitro* (Feng et al., 2018). Exogenous calcium and boron application, proposed to enhance pectin-pectin crosslinks, can complement, to some extent, FER-dependent salt-induced phenotypes suggesting that salt application might affect pectin architecture (Feng et al., 2018). No available information exists for a role of other CrRLK1L members in sensing salt induced cell wall damage. However, other cell wall localized proteins such as the Leucine-rich repeat extensins (LRX) 3/4/5 have been reported be involved in helping the physical association between RALF22/23 and FERONIA causing FER inactivation through its internalization upon salt stress (Zhao et al., 2018).

Here we report that salt changes pectin architecture, and that these modifications appear to reduce the degree of methyl esterification of the main component of pectin, homogalacturonan. We propose that this event is responsible for the activation of a range of early cellular signaling, gene expression and phenotypic responses. In our attempt to expand our current knowledge on the role of other CrRLK1L in the abiotic field, we have analyzed a complete set of salt stress responses to shed some lights on the insight of early and late salt stress signaling events in previously characterized mutants. We also show that while loss of function mutants lacking THE1 or HERK1 alone show wild type like salt stress responses, plants lacking both HERK1 in combination with a truncated, gain of function allele version of THESEUS1 (*herk1the1-4* mutants (Li et al., 2009)) show severe salt sensitivity, similarly to that of *fer-4* loss of function mutants. FER and HERK1/THE1-4 are required for dampening the salt-induced Mitogen Activated protein Kinase 6 (MPK6) activation, as higher activation of MPK6 is observed in *fer-4* and *herk1the1-4* mutants which is consistent with the higher expression of salt induced, cell wall damage dependent, marker genes in these mutants. We also report that the salt-induced pectin methyl esterification of loosely bound pectins can be rescued by the pectin methyl esterase inhibitor [(-)-Epigallocatechin gallate (EGCG)] or calcium chloride (CaCl_2_) application. Here we suggest that salt induced modifications are sensed by FER, HERK1/THE1-4 combination which are required to inhibit downstream responses partly dependent on MPK6 function. Collectively, we show that salt changes the degree of pectin methyl esterification, likely affecting the structural integrity and the mechanical properties of the cell wall, perceived by receptor-like kinases, which are in turn responsible for regulating a range of salt stress induced responses.

## Results

### herk1the1-4 and fer-4 mutants are affected in early responses to salt

To investigate whether other CrRLK1Ls, besides *fer-4,* are disturbed by salt application (Feng et al., 2018; Zhao et al., 2018), a collection of available mutants in members previously implicated in CWI sensing (Table S1), were directly germinated and grown in the presence/absence of 150 mM NaCl for 10 days (Figure S1A). No bleaching was observed in control plates (not shown) and the differences between the genotypes were only quantified in salt containing plates (Figure 1A). After 10 days of growth, *fer-4* mutants showed higher rates of cotyledon bleaching as expected, while both *herk1* and *the1-1* single knock-out (KO) mutants displayed comparable salt induced cotyledon bleaching to that observed in salt-grown wild type (wt) controls. On the other hand, slightly more bleaching was discovered in the single *the1-4* mutant that expresses a truncated THE1 protein, and was previously described as a gain of function allele of THE1 (Merz et al., 2017). Interestingly, the double *herk1the1-4* showed enhanced bleaching comparable to *fer-4* (Figure 1A), suggesting that a combination of HERK1 and THE1-4 is required to regulate salt-dependent sensitivity. Genetically combining *the1-1* loss of function and *fer-4* mutant led to the same bleaching phenotype as *fer-4* single mutants (Figure 1A/S1A) suggesting that the FER-dependent salt induced phenotype cannot be alleviated by impairing the functionality of the cell wall damage sensor THE1. To better investigate the role of CrRLK1Ls mutants in salt induced signaling, different experimental set-ups have been employed to analyze early (MPK6 phosphorylation, gene expression) and late responses (root halotropism, cotyledon bleaching and cell wall composition) to salt stress (Figure 1B).

**Figure 1.**
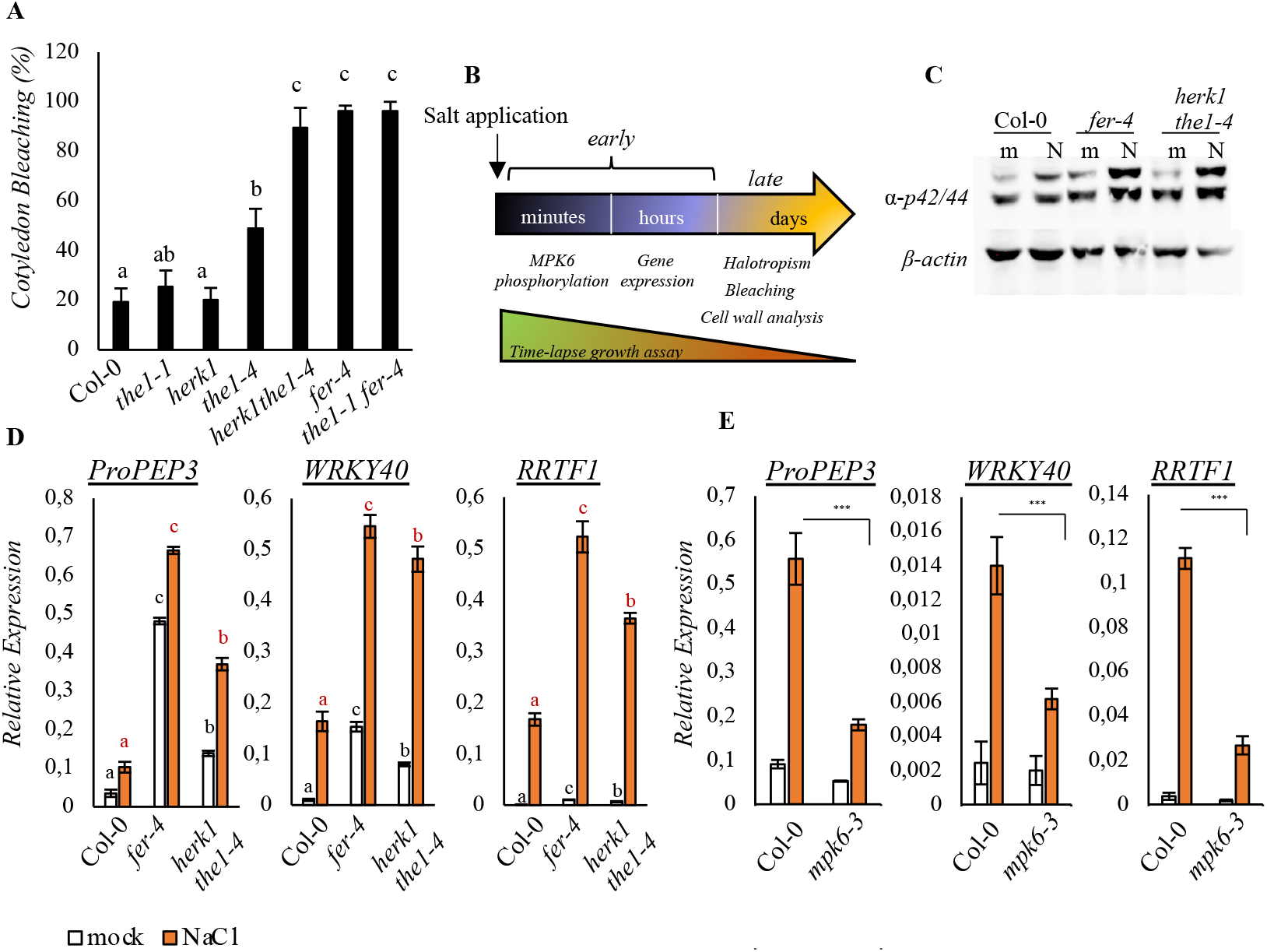
*herk1the1-4* and *fer-4* mutant lines show enhanced early responses to salt stress. **A.** Cotyledon bleaching analysis was conducted in seedlings grown for 10 days in the presence of 150 mM NaCl. Values, expressed in percentage are the average of 4 independent experiments. Bars represent Standard Error (SE). One-way ANOVA and Tukey’s HSD (*α*= 0.05) were used to compare differences between genotypes within one treatment condition (salt). **B.** Timeline of the treatments performed in this paper together with the terminology used. **C.** 7-day old seedlings of Col-0, *fer-4* and *herk1the1-4* were grown in liquid 0.5% MS and treated for 15 minutes with H_2_0 (mock, m) or 100 mM NaCl (N). Phosphorylation of MPKs was detected by immunoblot assay using specific antibody against the pMPK6 and pMPK3 (α-p44/42-ERK). Equal loading performed upon stripping was carried out by using anti *β*-actin Arabidopsis antibody. Similar results were obtained in 3 biological replicates (n= 3). **D.** Seedlings as in **C** were treated for 1h to analyze the expression levels of selected salt-induced marker genes. Expression of *ProPEP3, WRKY40, RRTF1* relative to *ACT2*, determined by qRT-PCR is represented in the histograms as the average of 3 biological replicates (n= 3). Bars indicate standard deviation (SD). Statistical comparisons have been made by using One-way ANOVA and Tukey’s HSD (*α*= 0.05) considering the 3 different genotypes within the same treatment (black letters for mock comparisons, red for NaCl treatments). **E.** 7-day old seedlings of Col-0, *mpk6-3* were treated as in D. and gene expression analysis was performed after 1h treatment H_2_0 (mock, m) or 100 mM NaCl. n= 3. Bars indicate standard deviation (SD). Asterisks show statistical comparisons between NaCl-treated Col-0 and NaCl-treated *mpk6-3* according to Student’s t test, ***P < 0.001).

To understand whether salt-triggered oversensitive phenotypes observed in *fer-4* as well as in *herk1the1-4* are the result of enhanced salt-dependent signaling we analyzed early responses upon salt application. In salt treated plants, one of the early responses is the phosphorylation of MPK6 (Yu et al., 2010). In plants and animals, MAPK cascades are initiated by a specific stimulus which triggers the activation of MAPK kinase kinases (MAPKKKs), phosphorylating a MAPK kinase (MAPKK) and in turn leading to the activation of MAPKs, also known, in Arabidopsis as MPKs (Morrison, 2012). By using a well characterized and widely used antibody (Bigeard and Hirt, 2018; Galletti et al., 2011; Savatin et al., 2014) that recognizes the phosphorylated form of both Arabidopsis MPK6 and 3, we have analyzed salt induced MPK6 phosphorylation upon 15 minutes of treatment. We first performed control experiments in a previously characterized MPK6 allele (*mpk6-3*) (Yu et al., 2010) to confirm the specificity of the antibody (Figure S1B). Then we used the same set-up to perform analysis in the mutants. Both *fer-4* and *herk1the1-4* mutant lines showed a stronger MPK6 activation when compared to NaCl-treated wt seedlings (Figure 1C/ S1C). Interestingly, similar to the bleaching phenotype, while *herk1* and *the1-1* did not show a difference with salt-treated wt control, *the1-4* displayed enhanced salt-induced MPK6 activation when compared to the wt, although less evident when compared to the corresponding double *herk1the1-4* mutants or *fer-4* (Figure S1C).

To investigate whether the lack of FER or HERK1/THE1-4 had an effect on salt-induced gene expression levels, we selected three salt-induced marker genes (Engelsdorf et al., 2018; Loo et al., 2020; Nakaminami et al., 2018; Soliman and Meyer, 2019) and checked their expression in response to salt application. We chose *ProPEP3, WRKY40* and *RRTF1* because they are strongly induced within 1 h from salt application and because they have been described responsive to a wide number of salt dependent modifications, like cell wall changes, ABA-dependent signaling and reactive oxygen species (ROS) (Engelsdorf et al., 2018; Loo et al., 2020; Nakaminami et al., 2018; Soliman and Meyer, 2019) (Table S2). We treated wt, *fer-4* and *herk1the1-4* seedlings for 1 h and found that the expression level of salt responsive genes was higher in response to salt in the mutant lines when compared to the wt controls (Figure 1D). Similar to the cotyledon bleaching rates, *the1-4* but not *the1-1,* showed enhanced transcriptional regulation of salt marker genes compared to salt treated wt seedlings (Figure S1D).

Since *mpk6-3* loss of function mutants were reported to show reduced root growth inhibition in response to salt stress (Han et al., 2014), likely associated to reduced salt sensitivity, we analyzed the expression pattern of the selected gene markers in the *mpk6-3* mutant background and observed that MPK6 is at least in part responsible for the upregulation of *PROPEP3, RRTF1* and *WRKY40* by salt (Figure 1E). The results suggest that the enhanced MPK6 activation observed in *fer-4* and *herk1the1-4* might also be responsible for their amplified susceptibility to salt. Recently it was reported that ectopic over expression of *RRTF1* is detrimental to plant salt tolerance (Soliman and Meyer, 2019). This is consistent with our observations, where the higher *RRTF1* expression pattern described in both salt-treated *fer-4* or *herk1the1-4* is linked to the higher plant salt susceptibility (Figure 1D/S1D).

### herk1the1-4 show a fer-4-like root growth inhibition dynamic after salt application

Next, because *fer-4* was previously reported to show defective root growth dynamics upon salt (Feng et al., 2018), we grew double mutant *herk1the1-4* seedlings in vertical plates and transferred them 7 days after in plates containing homogeneous NaCl (or no NaCl (control). To have a complete overview of both salt-triggered early and late effects, salt induced root growth inhibition was traced over 42 h in a specialized growth chamber where time lapse photography-based analyses were performed over time (Figure 2A). Previously, it has been reported that wt seedlings subjected to salt stress display a “multi-phasic” trend of growth, which is slowly inhibited within hours from initial salt application (quiescent phase) followed by a recovery phase (Geng et al., 2013). When compared to the wt, *herk1the1-4* mutant seedlings show, similar as reported for *fer-4* (Feng et al., 2018), reduced root elongation rate upon salt application associated with an impaired recovery upon the induction of the quiescent phase (Figure 2A).

**Figure 2.**
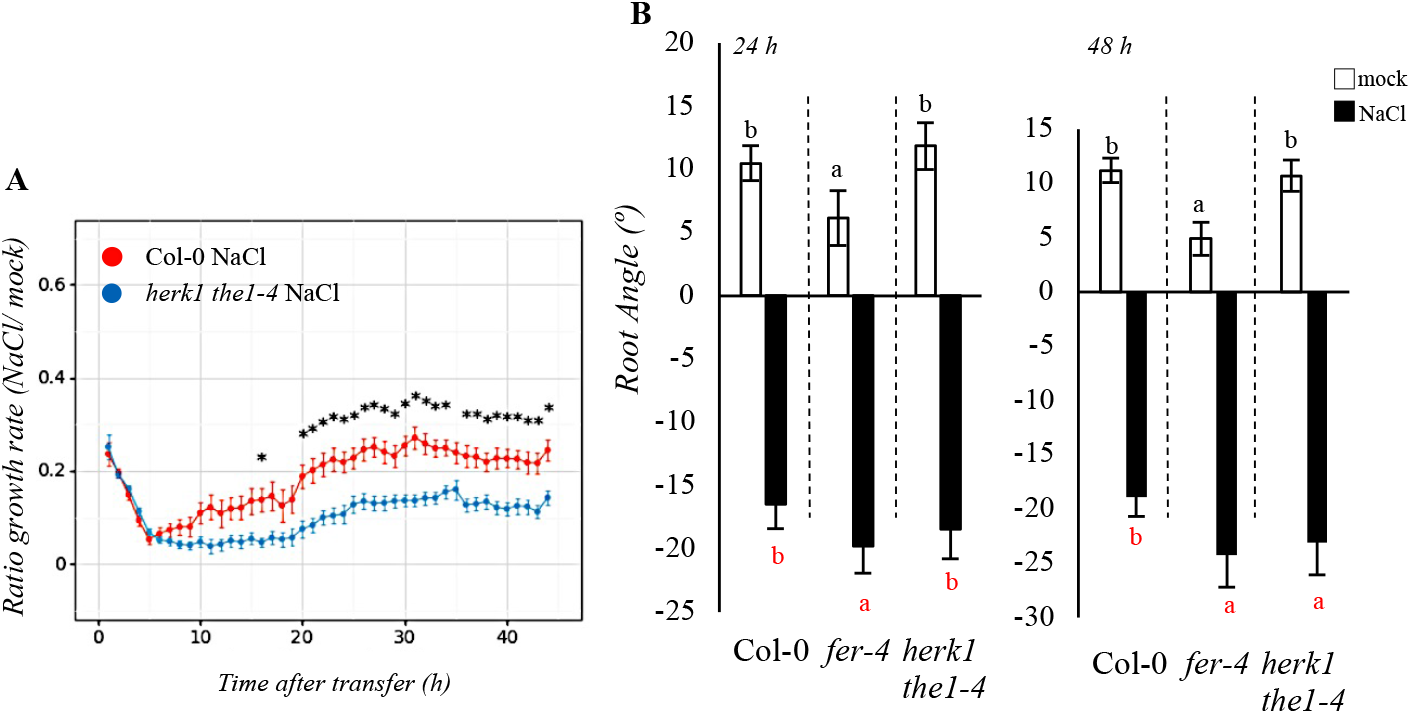
*herk1the1-4* and *fer-4* mutant lines show enhanced late responses to salt stress. **A.** Root growth rate was monitored for 42 h in a time lapsed growth chamber setting in 7-day old Col-0 and *herk1the1-4* transferred at time 0 h to new plates containing mock or 125 mM NaCl. Root elongation rate was calculated over 2 h intervals, averaged per hour and expressed as a ratio between NaCl and control treatment (mock). Each of the curve dots represent the average length and the corresponding standard error analyzed in three independent experiments (n= 45 seedlings per treatment, per genotype). Curves show the results of Statistical comparisons were performed by analyzing every timepoints for equal variance using a Levene’s test, followed by T-test to compare different genotypes at the same timepoint. The significance level (p= 0.05) was corrected for multiple testing by using stepwise Sidak correction (* = significant). **B.** 4-day old seedlings of Col-0, *fer-4* and *herk1the1-4* were subjected to NaCl gradient (for detailed explanation see Materials and Methods section) and root angle was analyzed with Smartroot 2.0 after 24 h or 48 h treatment. Histogram represents the average root angle of 3 independent experiments, bars show the average of standard errors (SE). One-way ANOVA and Tukey’s HSD (*α*= 0.05) were used to compare differences between genotypes within one treatment condition. Black letters show the comparisons between different genotypes in mock conditions while the red ones show those performed in the presence of NaCl. (n= 150).

To evaluate if HERK1/THE1-4 as well as FER regulate also other salt responses, we analyzed later responses such as halotropism and salt tolerance. When facing a salt gradient, plant roots bend to avoid high salt concentrations, a phenotype referred to as halotropism (Deolu-Ajayi et al., 2019; Galvan-Ampudia et al., 2013; van den Berg et al., 2016). Wild type, *fer-4* and *herk1the1-4* double mutant seedlings were subjected to diagonal (mock or salt) gradients and root bending [(angle) º] was measured 24 h and 48 h after salt application. When compared to the wt, *fer-4* exhibited slightly enhanced halotropism-triggered root bending after 24 h while *herk1the1-4* only displayed differences after 48 h [Figure 2B (individual biological replicates/root elongation rates shown in Figure S2A/B)]. The salt dependent root halotropism responses detected in *mpk6-3* [and shown as result of three independent experiments (Figure S2C)] display no significant differences when compared to the wt controls, suggesting that the mild halotropism defects observed in *fer-4* and *herk1the1-4* mutants are MPK6-independent. To evaluate whether the higher salt sensitivity observed in the double *herk1the1-4* was also present in adult plants we investigated the responses to salt in soil-grown plants (Figure S2D). For this assay, Col-0, *fer-4* and *herk1the1-4* plants were once watered with MilliQ water or NaCl (75 mM) and then rosette fresh weight was measured at the end of the treatment. Fresh weight quantification showed a significant difference between salt treated wt and the two mutant lines (Figure S2E) while reduced biomass was also observed in the control conditions (Figure S2F). Taken together, our results show that the *herk1the1-4* mutant exhibits a similar enhanced salt sensitivity as that detected in *fer* mutants not only for early stress responses (Figure 1), but also in growth dynamics and root elongation upon salt stress application (Fig 2).

### Salt-induced gene expression and growth responses, but not halotropism, are mitigated by addition of calcium

The effect of calcium in rescuing *fer-4* dependent salt induced root growth phenotypes was suggested to be associated to calcium mediated pectin crosslinking, and FER’s malectin domain was reported to bind pectins *in vitro* (Feng et al., 2018). To assess the effect of Ca^2+^ on signature responses to salt we treated seedlings with salt alone or in combination with 5 mM CaCl_2_. First, to understand whether calcium can influence the growth dynamics and intensity of the salt-triggered quiescent phase, we performed a time course analysis on wt seedlings grown in vertical conditions on control media for 7 days before being transferred to salt-containing plates. We observed that, as expected, salt induces an early drop in root elongation for the first hours upon salt application followed by a recovery phase that is fully completed after 20 h from the application of the stimulus. Compared to salt treatment, wt seedlings treated with both calcium and salt displayed a shorter and less severe quiescent phase and an early recovery phase (Figure 3A), while no visible effect on root growth was observed in the presence of calcium when compared to the mock control (Figure S3A).

**Figure 3.**
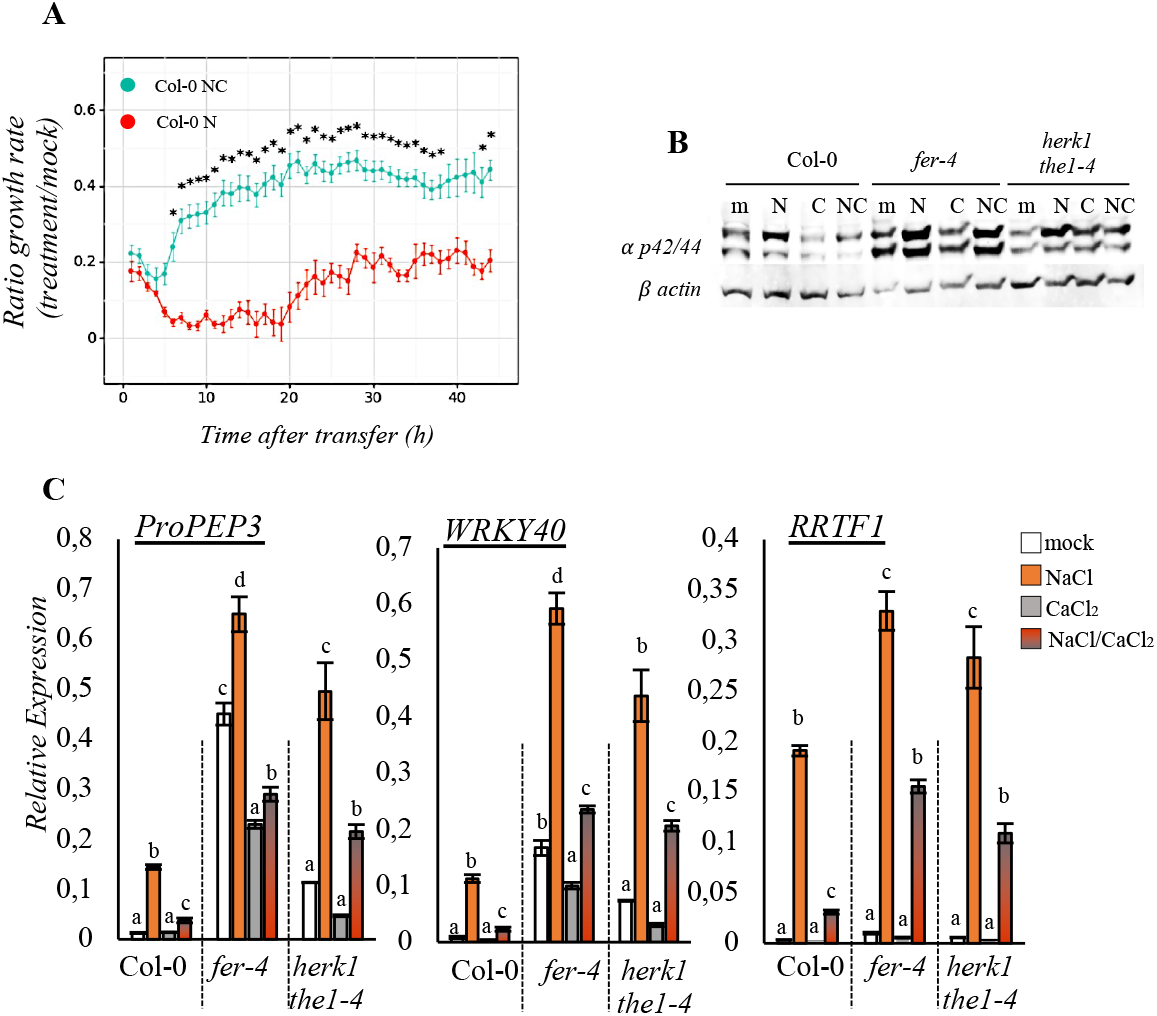
Salt-induced responses are alleviated by CaCl_2_ application. **A.** Root growth rate elongation monitored for 42 h in 7-day old Col-0 seedlings treated from 0 h (transferring time) with mock, 125 mM NaCl (N), 5 mM CaCl_2_ or a combination of NaCl+CaCl_2_ (NC). Root growth rate is expressed as a ratio between NaCl and mock treatment or NaCl+CaCl_2_ and mock. Each of the dots represent the average length and the corresponding standard error analyzed in two independent experiments (n= 30 seedlings per treatment). Statistical comparisons were performed by analyzing every timepoint for equal variance using a Levene’s test, followed by T-test to compare different treatments at the same timepoint. Samples were corrected for multiple testing by using stepwise Sidak correction. *, P< 0.05 **B.** 7-day old seedlings of Col-0, *fer-4* and *herk1the1-4* were grown in liquid 0.5%MS and treated for 15 minutes with H_2_0 (mock, m), 100 mM NaCl (N), 5 mM CaCl_2_ (C) or a combination of NaCl and CaCl_2_ (NC). MPKs phosphorylation detected by immunoblot assay, was performed by using specific antibody against the phosphorylated forms of both MPK3 and MPK6 (α-p44/42-ERK). Equal loading was carried out by using anti *β*-actin Arabidopsis antibody. Similar results are shown in figure S3B. **C.** Expression of *ProPEP3, WRKY40, RRTF1* determined by qRT-PCR and relative to *ACT2,* was performed in 7-day old seedlings of Col-0, fer-4, *herk1the1-4* treated as in B. for 1h. n= 3. Bars indicate standard deviation (SD). One-way ANOVA and Tukey’s HSD (*α*= 0.05) was used to analyze statistical differences between treatments within the same genotypes.

When analyzing early salt induced signaling response, we found that already after 15 minutes calcium could alleviate salt triggered MPK6 phosphorylation in the wt as well as in the salt oversensitive mutants *fer-4, herk1the1-4* (Figure 3B and S3B). Also, when wt, *fer-4* and *herk1the1-4* mutant lines were treated with a NaCl/CaCl_2_ combination, gene expression level was rescued to the same (*PROPEP3, WRKY40*) or almost similar (*RRTF1*) physiological expression levels when compared to the mock controls (NaCl/CaCl_2_ vs mock) (Figure 3C). As previously reported, sodium dependent root halotropism cannot be mitigated by exogenous calcium application (Figure S3C/D, and (Galvan-Ampudia et al., 2013)). Thus, while Ca^2+^ has been suggested before to function as a cell wall strengthening agent which can mitigate salt induced *fer-4* cell swelling phenotypes, we now show that its effects are likely to happen earlier in time not only on a structural, but also on a cellular signaling level.

### Salt induced homogalacturonan de-methyl esterification is partly required to activate salt-triggered responses

In cell walls, calcium ions have been connected to enhanced wall strength, and described to play a role in the formation of calcium ion-mediated homogalacturonan-homogalacturonan (HG, the main pectin component) crosslinks (Domozych et al., 2014). For this reason, the hypotheses for calcium being responsible to alleviate salt induced responses have been associated to its ability to alter pectin architecture. To study salt induced cell wall modifications directly, we analyzed pectin isolated from 6-day old wt seedlings treated with or without salt for 24 h. The extracted cell wall material was subjected to pectin extraction by using boiling distilled water (see Material and Methods). Total sugars in these extracts were quantified (Dubois et al., 1951; DuBois et al., 1956) and sugars were spotted on nitrocellulose membrane to perform dot blot analyses. To examine in detail the structural organization of pectins and in particular the degree of methylation (DM) of HG we used three cell wall probes, namely 2F4, LM19 and LM20 which respectively recognize partially de-methyl-esterified, fully de-methyl-esterified and highly methyl-esterified HG (Peaucelle et al., 2008; Verhertbruggen et al., 2009) (Figure S4A). As a control, HG methyl esters were removed by incubating the same amount of total sugars spotted in the upper part of the dot blot panels (native pectins) with high pH buffer (Na2CO3, pH 11, (deM-HG)) providing loading controls for both 2F4 and LM19 and negative controls for LM20 specificity. 2F4, a probe previously reported to specifically recognize Ca^2+^-crosslinked HG dimers (Douchiche et al., 2010; Liners et al., 1989) shows enhanced signals in the Na2CO3 treated extracted sugars (Figure 4A, first lower panel), suggesting that this probe in fact binds partly or fully de-methyl esterified HG [as also reported in (Peaucelle et al., 2008)]. When wt seedlings were treated with salt, we observed that water-extracted cell wall pectins displayed enhanced 2F4 and slightly enhanced LM19 signal, while no differences were detected in LM20 incubated samples, revealing an effect for salt in altering pectin methyl esterification patterns by decreasing the concentration of methyl esters on the homogalacturonan (Figure 4A). Analysis of pectin methyl esterification in the salt sensitive *fer-4* and *herk1the1-4* mutants showed that similarly to the wt they also display changes in the overall methyl esterification distribution upon salt stress suggesting that in these mutants, structural modifications of the cell walls were also induced by salt treatments similar to wt but were not detectable in mock conditions (Figure 4B/S4B).

**Figure 4.**
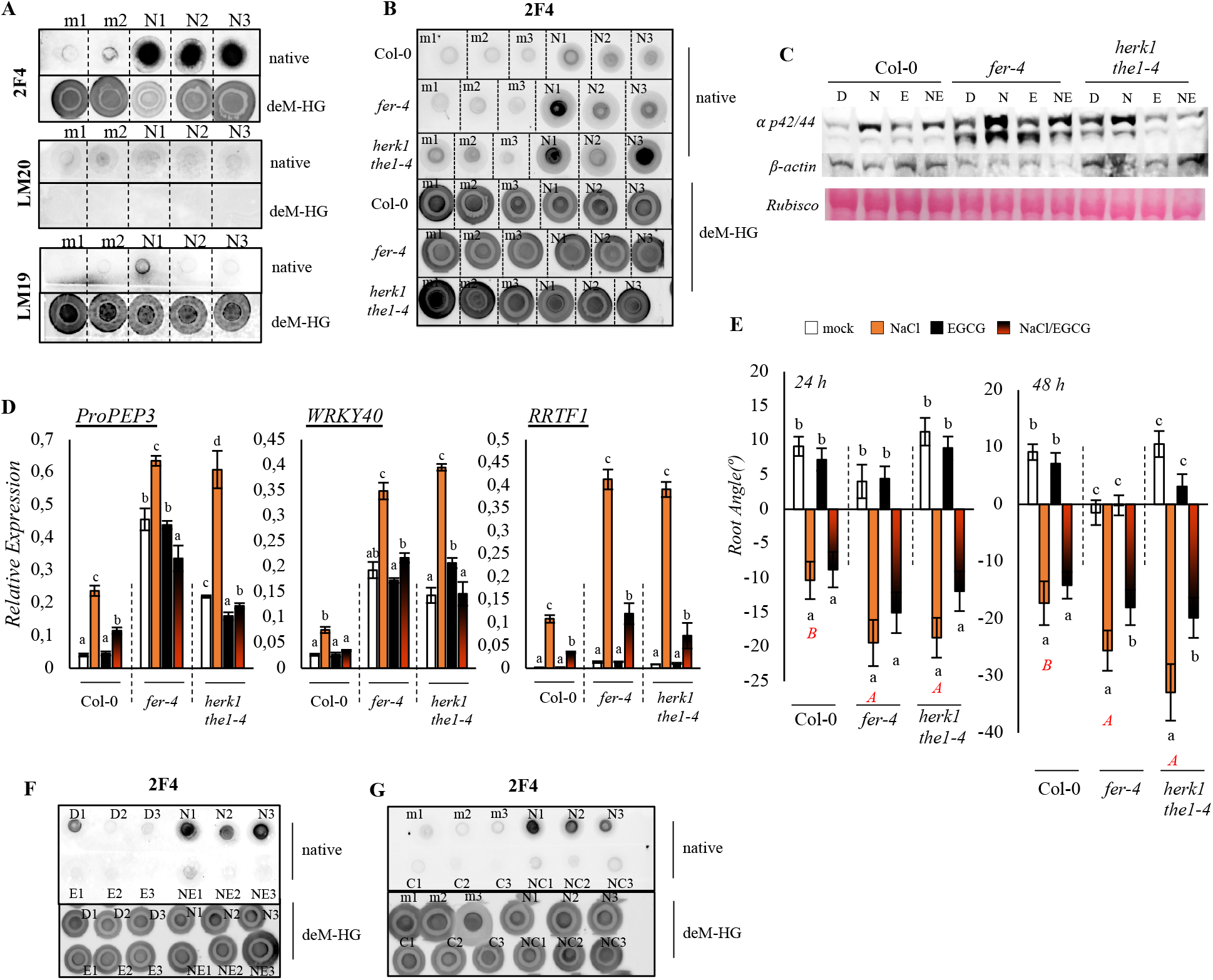
Salt induced alteration methylation of loosely bound pectins can be alleviated by CaCl_2_ or EGCG application. **A.** Six-day old Arabidopsis wild type seedlings were grown in flasks and treated for 24 h with H_2_0 (m) or 100 mM NaCl (N). 3 biological replicates, containing 50 seedlings each, for mock and NaCl were used to extract the Alcohol Insoluble Residues (AIRs). Pectin factions were obtained by boiling the AIRs with distilled sterile water (pH 7.2) as described in materials and methods. The extracted sugars were quantified and 5 *μ*g of total sugars were transferred in tubes containing the same volume of ddH_2_O (native) or 0.1M final concentration of Na_2_CO_3_ pH 11, to remove methyl esterification [demethylated HG (labeled with deM-HG)]. After 1 h incubation, sugars were spotted on nitrocellulose membranes. Dot-blots were performed with 2F4 (in T/C/N buffer) or LM20 and LM19 (in PBS+BSA buffers). **B.** Six-day old seedlings of Col-0, *fer-4* and *herk1the1-4* mutants were grown and treated as in A. Sugars were extracted as in A and 5 *μ*g spotted on nitrocellulose membranes after 1 h incubation with ddH_2_O (native) or Na_2_CO_3_ pH 11 (deM-HG). Dotblot was performed with 2F4**. C.** 7-day old seedlings of Col-0, *fer-4* and *herk1the1-4* were treated for 15 min with DMSO (D), 100 mM NaCl (N), 50 *μ*M EGCG (E) and NaCl/EGCG (NE). 15 *μ*g of total proteins loaded and phosphorylation of MPK6 analyzed via western blot assays with *α* p42/44 antibody. β-actin and Ponceau Red are shown for equal loading. **D.** Expression of *ProPEP3, WRKY40, RRTF1* determined by qRT-PCR and relative to *ACT2*, was performed in 6-day old seedlings of Col-0, *fer-4, herk1the1-4* treated for 1 h with DMSO (mock), 100 mM NaCl, 50 *μ*M EGCG and NaCl/EGCG. n=3. Bars indicate standard deviation (SD). Statistical comparisons have been made by using One-way ANOVA and Tukey’s HSD (*α*= 0.05) considering the 3 different genotypes within the same treatment (black letters for mock comparisons, red for NaCl treatments). **E.** 4-day old seedlings of Col-0, *fer-4* and *herk1the1-4* were subjected to halotropism assay with DMSO (mock), 100 mM NaCl, 50 *μ*M EGCG and NaCl/EGCG gradients. Root angle was after analyzed after 24 h, 48 h and 72 h. Histogram represents the average root angle of 3 independent experiments, while bars show the standard errors (SE). (n= 75). Letters display One-way ANOVA and Tukey’s HSD (*α*= 0.05) within the same genotype. Red capital letters represent statistically significant differences according to one-way ANOVA and Tukey’s HSD (*α*= 0.05) between genotypes within one treatment condition. **F.** Six-day old Arabidopsis wild type seedlings were grown in flasks and treated for 24 h with DMSO (D), 100 mM NaCl (N), 50 *μ*M EGCG (E) and NaCl/EGCG (NE). (n= 3). **G.** Similarly, 3 other biological replicates, were treated with H_2_0 (m) or 100 mM NaCl (N), 5 mM CaCl_2_ (C) or a combination of NaCl and CaCl_2_ (NC). As in G 5 μg of total sugars were transferred in tubes containing the same volume of ddH_2_O (native) or 0.1M final concentration of Na2CO3 pH 11, to remove methyl esterification (deM-HG). Dot-blot was performed with 2F4.

To investigate the cellular and physiological consequences of salt induced HG de-methyl esterification, a chemical PME inhibitor, (-)-Epigallocatechin gallate (EGCG), was used to perturbate PME activity (Wolf et al., 2012). Therefore, seedlings of wt, *fer-4* and *herk1the1-4* were treated with a combination of salt and ECGC and effects were compared to the mock (DMSO; EGCG solvent). We found that even if ECGC alone did not affect MPK6 activation in any of the genotypes, when the combination NaCl/EGCG was applied, MPK6 activation was reduced especially in the salt sensitive mutants (Figure 4C, S4C), suggesting that inhibition of salt-induced de-methyl-esterification of pectin can alleviate the immediate stress responses reported in this study. Similarly, to determine the effect of EGCG on salt-induced gene expression levels, seedlings of Col-0, *fer-4* and *herk1the1-4* mutants were treated with NaCl, EGCG or NaCl/EGCG. During these experiments we observed a similar trend as in seedlings treated with CaCl_2_/NaCl combinations. While EGCG itself did not affect the expression of the selected markers, when EGCG was applied in combination with NaCl, gene expression induction levels were reduced most evidently in the salt sensitive *fer-4* and *herk1the1-4* mutants (Figure 4D). To study the effect of EGCG on root halotropic response, we analyzed root direction (angle) for seedlings treated with NaCl on a gradient supplemented with or without EGCG. A mild but significantly reduced root bending was detectable in the *herk1the1-4* and *fer-4* mutants, but not Col-0 seedlings, after 48 h of co-treatment (NaCl+EGCG) [(Figure 4E), biological replicates and root elongation rates are shown in Figure S4D/S4E/S4F]. EGCG itself did not alter root direction at any time point when compared to the mock control (Figure 4E, S4D/S4E, black vs white bars).

To analyze the effect of EGCG on salt-induced pectin de-methyl esterification, we analyzed HG methylation patterns and discovered that when both salt and EGCG were present the amount of de-methyl esterified pectin recognized by 2F4 (Figure 4F) was reduced compared to that detected in seedlings treated with salt only, while as expected, LM20 signal was not affected (Figure S4G). Interestingly, when a similar experiment was performed where wt seedlings were treated with a combination of salt and calcium (Figure 4G), calcium was also found to attenuate the 2F4 dependent signals, suggesting that salt-dependent de-methylation of HG might lead to reduced cell wall strength which can be reinforced by exogenous application of calcium, and/or that calcium would work indirectly to affect the regulation of enzymes required for the salt induced pectin relaxation. Yet, these hypotheses remain to be addressed.

Taken together these results suggest that salt-induced modification at the cell wall level involves pectin architecture alterations and that PME inhibition can partially counteract protein kinase phosphorylation and salt dependent gene expression in salt sensitive genotypes. Even if our cell wall analyses were performed 24 h after salt application, it is possible that changes in pectin moieties happen quickly at the single cell level. However, by using our method, these modifications become only detectable later upon the treatment, when the effect on pectin is widely spread in all cell types. This hypothesis would explain why salt induced intracellular responses can be attenuated by Ca^2+^/EGCG application.

Collectively our data show that a large part of the salt induced responses are triggered after perception of salt induced pectin architecture modification. Both calcium and EGCG application alleviate not only the salt induced pectin de-methyl esterification but also downstream responses induced by salt, such as MPK6 phosphorylation and gene expression, which appear to be negatively regulated by the cell wall sensors FER and HERK1/THE1-4 combination.

## Discussion

Cell walls are complex polysaccharide networks composed of cellulose, hemicelluloses, pectins and structural proteins (Höfte and Voxeur, 2017). Pectins are abundant polysaccharides in Arabidopsis primary cell walls, that are rich in galacturonic acid and create homogalacturonan (HG) chains made of linear α-1-4-linked galacturonic acid monomers. HG is transported to the cell wall in a highly methylated form and de-methyl esterified in the apoplast by the action of pectin methyl esterases (PMEs) (Lampugnani et al., 2018). It has been suggested that de-methyl esterified HG can be crosslinked by Ca^2+^ and form so-called egg-boxes, pectin crosslinked structures involved in promoting cell wall stiffening (Hocq et al., 2017). However, the hypothesis that calcium crosslinks limit cell growth is not well supported, as the analysis of calcium mediated pectin crosslinks is challenging (Wang et al., 2020). In this study we report that while external calcium application does not influence root growth responses, in combination with salt, calcium enhances root growth likely limiting the inhibitory effects that salt has on cell elongation (Figure 3A). It has been reported that pectin modifications and crosslinking influence the strength of the wall as well as cell shapes (Pelloux et al., 2007; Rui and Dinneny, 2020). Yet, while in pollen tubes the data fit the hypothesis that lower HG esterification is associated with higher cell wall stiffening, several other studies show that reduction of HG methyl esterification patterns rather enhance cell wall expansion (Haas et al., 2020; Peaucelle et al., 2011). Analysis performed by atomic force microscope (AFM)-based indentation on shoot apical meristems of plants with reduced pectin methyl esterification, showed enhanced cell wall softening (Haas et al., 2020; Peaucelle et al., 2011). It has been hypothesized that in the absence of calcium, demethyl esterified pectins displayed altered fluidity, being more susceptible to the action of pectin degrading enzymes like polygalacturonanses (PGs) which enhance cell wall extensibility (Peaucelle et al., 2008; Wang et al., 2020).

In recent years a mechanism required to monitor cell wall structure named cell wall integrity (CWI) maintenance has been described in plants. CWI is essential for maintaining cell wall plasticity and initiating responses to cell wall damage (CWD), which can be triggered by both biotic or abiotic stress (Vaahtera et al., 2019). Cell wall damage sensor candidates include receptor-like kinases (RLKs), Leucin-rich repeat extensins (LRXs) and ion channels [reviewed in (Rui and Dinneny, 2020)]. Well-described CWI sensors belong to the *Catharanthus roseus* Receptor-Like Kinase1-Like (CrRLK1L) family, such as THESEUS1 (THE1) and FERONIA (FER) (Cheung and Wu, 2011). Previous research has shown the essential role of THE1 in mediating the response to cellulose impairment and suggested its function as a cell wall damage sensor (Engelsdorf et al., 2018; Hématy et al., 2007). Several mutant lines for THE1 have been reported to date. The THE1 loss of function mutant (*the1-1*) is almost insensitive to treatment with the cellulose biosynthesis inhibitor isoxaben (ISX), while *the1-4* is a gain of function allele, lacking the intracellular kinase domain, which displays higher ISX susceptibility (Engelsdorf et al., 2018), highlighting its essential function in sensing CWI changes. In our attempt to better understand the role of CrRLK1Ls during abiotic stress and in particular in regulating salt stress responses, a collection of mutants previously described for their role in CWI maintenance has been characterized. Amongst them, we included the double *herk1the1-4* mutants originally reported lacking the function of both the CrRLK1Ls THE1 and HERKULES1 and only later characterized for being a combination of *herk1* loss of function and *the1-4* (Merz et al., 2017). These double mutants share similar stunted growth phenotypes with *fer-4* adult plants, while loss of function mutations for HERK1, or THE1-4 show a wild type-like physiology in terms of cell elongation and rosette leaf shape (Li et al., 2009). In our experimental conditions, we observed that *herk1-the1-4* mutant lines not only shared similar vegetative growth but also displayed a *fer-4-like* enhanced salt sensitivity. It is interesting to notice that in the presence of salt even if *the1-1* showed no significant differences when compared to the wild type controls (as also reported in (Van der Does et al., 2017)), the single *the1-4* gain of function allele also showed mildly enhanced salt induced responses. The *herk1the1-4* double mutants were even more disturbed than the corresponding single mutants suggesting that combinations between these two CrRLK1Ls is required for mimicking *fer-4* in salt and non-salt-induced phenotypes.

To date, potential transient hetero-oligomerization of multiple CrRLK1Ls has been suggested, for example in the case of HERK1/ANJEA or ANX/BUPS in pollen tubes [reviewed in (Ge et al., 2019)], thus a possible interaction between HERK1/THE1-4 regulating the activity of FER cannot be excluded. FER has been reported to function as scaffold and regulate the interaction with other plasma membrane localized proteins being responsible for modulating immune responses and suggesting a complex role for FER as a signaling hub intersecting several pathways and playing multiple functions in regulating plant growth/development and reproduction (Chen et al., 2016; Dünser et al., 2019; Stegmann et al., 2017). *fer-4* mutants exhibit significant growth defects and their root cells burst in response to salt application (Feng et al., 2018), but to date no available data have connected other CrRLK1Ls, besides FER, to salt-induced responses. The observed phenotypes have been connected with the hypothesis that FER directly senses cell wall pectin perturbations proposed to likely happen when Na^+^ ions interfere with pectin crosslinks (Feng et al., 2018) and FER’s malectin domain has been suggested to bind specifically de-methyl esterified pectin domains (Lin et al., 2018). Yet, all the available information is from *in vitro* interactions. Similarly, in yeast, the extracellular domain of the cell wall integrity WSC1 receptor, required to activate responses triggered by plasma membrane/cell wall displacement during osmotic stress, can bind to structural elements of yeast cell walls. WSC1, which is connected to the perception of mechanical stress and not to polysaccharide perception, functions as a nanospring involved in sensing the osmo-dependent cell wall/plasma membrane interphase modification upon cell shrinking or swelling (Dupres et al., 2009; Heinisch et al., 2010). In plants, THE1 has been hypothesized to function in a similar manner as the yeast WSC1, being required to activate responses induced by cellulose impairment-induced cell swelling. Interestingly ISX-induced responses could be alleviated by osmotic treatments which can rescue the cell wall/plasma membrane displacement to the control conditions (Hamann et al., 2009).

Here, we report that changes in cell wall elasticity observed in response to salt (Feng et al., 2018) might be the result of modifications in the degree of methyl-esterification of homogalacturonan, since wt seedlings treated with salt showed enhanced de-methyl esterified epitopes of pectin when compared to the mock controls (Figure 4A). It has been previously suggested that Na^+^ ions can intercalate in the de-methyl esterified pectin limiting calcium mediated pectin crosslinks. Therefore, excessive calcium ions can counteract the effect of sodium rescuing the right balance of crosslinked pectins (Feng et al., 2018). However, in our experiments we have observed that only in the presence of salt the reduction in methyl esters becomes visible, likely suggesting that salt might enhance the activity of enzymes required for the establishment of the pectin moieties. In our study we were able to detect cell wall changes at the whole seedling levels only after 24 h, but we hypothesize that small local (non-detectable) cell wall changes might happen immediately after salt application since salt-induced responses were alleviated by exogenous application of chemicals that alter pectin structure [such as calcium chloride and the pectin methyl esterase (PME) inhibitor ECGC (Dünser et al., 2019; Wolf et al., 2012) (Figure 4F/4G)]. This suggests that the local salt-induced pectin de-methyl esterification is sensed/perceived via the cell wall sensors FER alone or the combination of HERK1/THE, being in turn responsible for activating several main intracellular signaling pathways in plants. We also observed that while exogenous calcium can mitigate the salt-dependent HG de-methyl esterification as well as salt-triggered immediate/early responses it cannot alter the sodium specific root halotropic responses. These results are in accordance with (Galvan-Ampudia et al., 2013), where external calcium application did not change the degree of root salt avoidance.

In our experimental conditions, the salt sensitive *fer-4* and *herkthe1-4* mutant lines showed a mild but enhanced salt induced root bending after 48 h from salt application. Root halotropism has been reported to be independent of extracellular calcium and regulated by PIN-FORMED2 (PIN2) re-localization (Galvan-Ampudia et al., 2013) and seems to be independent on the function of MPK6 (Figure S2C). Whether the slightly enhanced halotropic response in salt-challenged *fer-4* and *herk1the1-4* could be due to mis-regulation of PIN re-localization in response to salt stress is yet unknown. However, in a recent paper it has been reported that the physiological root waviness phenotypes observed in *fer-4* seedlings grown in tilted plates depends on altered PIN2 localization (Dong et al., 2019), which might also explain our observations. We also observed that EGCG could alleviate the exaggerated salt induced root bending phenotypes observed in *fer-4* and *herkthe1-4* mutant lines. While the molecular mechanism is still unclear, it has been reported that EGCG application can affect the localization and dynamics of several plasma membrane localized proteins (including PIN3 and FLS2) (McKenna et al., 2019), likely suggesting that the ECGC dependent alteration of protein dynamics could impair the functionality of proteins required for the halotropic stress responses in salt sensitive mutants.

On the other hand, both salt-triggered MPK6 activation and gene induction can be mitigated by co-treatments with calcium or EGCG, suggesting that these salt stress responses are activated upon sensing the salt-dependent pectin moieties modification. Whether calcium application can counteract salt induced responses by inhibiting salt dependent PME activation could be hypothesized. Experiments performed to enhance tomato shelf-life, showed that fruits treated with CaCl_2_ show less PME activity and have enhanced tissue firmness (Mansourbahmani et al., 2017). In literature the role of PMEs and PME inhibitors (PMEIs) in regulating salt stress responses is far from being understood. For example, it has been reported that reduced expression of *PMEI10 (AT1G62760*) led to reduced salt sensitivity, likely suggesting that at least PMEI10 might function as negative regulator of salt stress responses [(Jithesh et al., 2012), also reviewed in (Wormit and Usadel, 2018)]. On the other hand, overexpression of *PMEI13* led to enhanced salt tolerance (Chen et al., 2018) fitting with our current observations, where PME mediated-salt-induced HG de-methyl esterification is required to activate at least in part salt-dependent signaling responses. This hypothesis is supported by our observations that salt induced cell wall damage-triggered responses are alleviated by chemicals that function in regulating pectin network organization, especially in *fer-4/herk1the1-4* mutants. Thus, a model could be hypothesized (Figure 5) where salt induces pectin de-methylation through PMEs, which is sensed by FER and HERK1/THE1-4.

**Figure 5.**
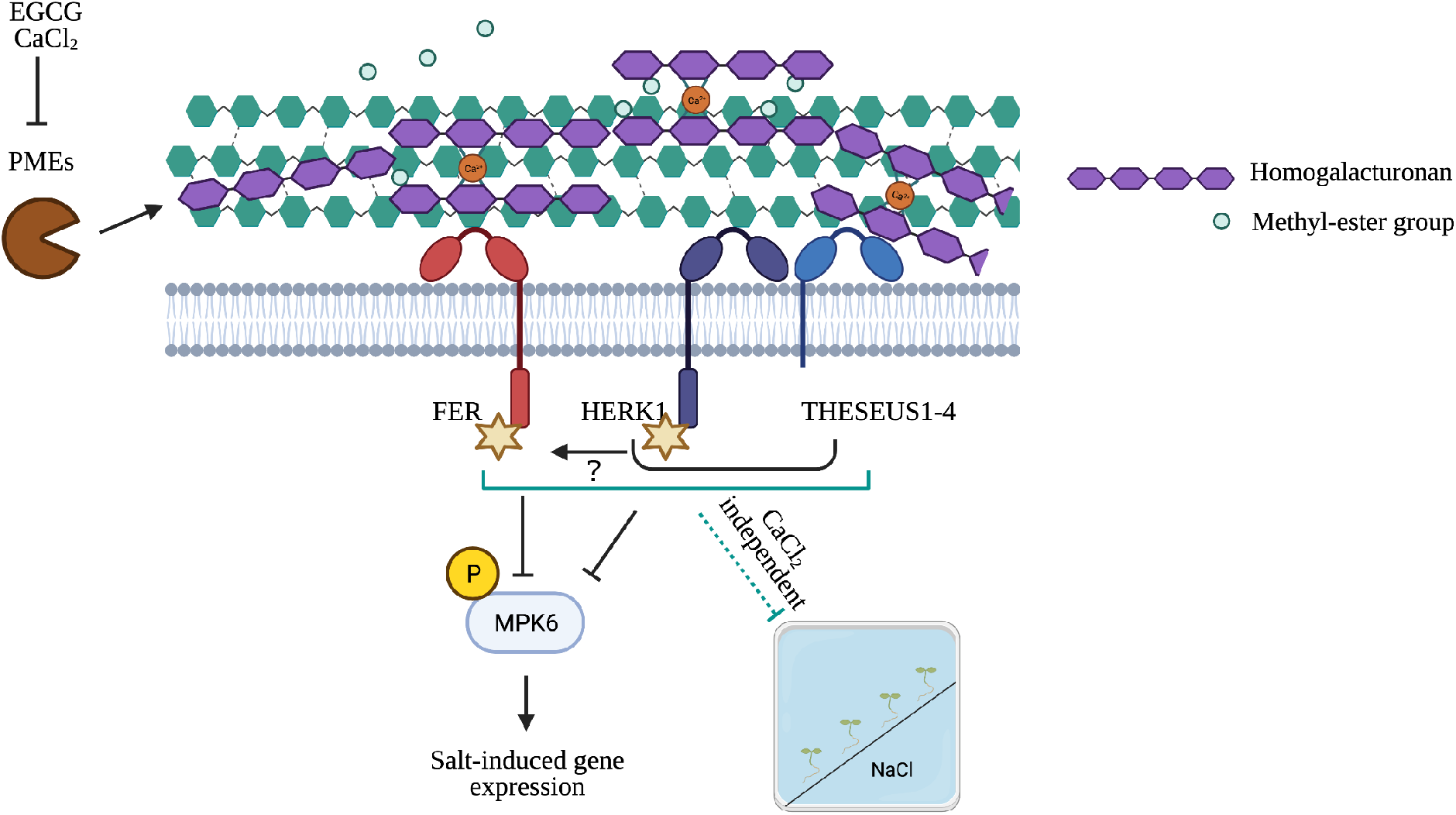
Hypothetical model based on the findings of this study. Salt treatment alters the degree of methyl esterification of homogalacturonan causing cell wall damage induced intracellular responses. Application of exogenous calcium or PME inhibition, can alleviate cell wall alterations likely mitigating intracellular signals. The cell wall sensor FERONIA (FER) is responsible for attenuating most of the salt dependent phenotypes is mimicked by HERKULES1 (HERK1) and a truncated version of THE1 (namely THE1-4). We hypothesize that the cell wall sensors HERK1 and THE1-4 might interact together to maintain FER functionality. Salt induced pectin modification has been observed in the wt as well as in these mutants suggesting that these sensors might be only functioning as antennas to perceive the damage. The regulation of salt dependent root bending known to require PIN2 re-localization seems to be mildly altered in plants lacking FER or HERK1/THE1-4 combination but seems to be independent on MPK6 function which is instead required for the full expression of salt induced marker genes.

## Conclusions

In this study we report and characterize in detail early and late salt stress-induced responses of seedlings (Figure 1B; Figure 5). We describe that salt induced pectin de-methyl esterification in the cell wall is perceived not only by the cell wall sensor FERONIA (FER) alone but also by the HERKULES1 (HERK1)/THE1-4 gain of function combination, exhibiting a similar role in salt stress related downstream signaling and responses. Pectin modification is required for the activation of downstream responses, such as MPK6 phosphorylation, which is in turn required for the full activation of salt-induced marker gene expression. Inhibition of MPK6 activation was described before to make plants more resistant to salt (*e.g.* enhanced root growth, reduced salt-triggered Ca^2+^ spike accumulation) (Han et al., 2014). In accordance, we show that MPK6 negatively regulates a major part of the cell wall modification induced salt stress responses (salt induced gene expression) which could affect salt tolerance. The higher MPK6 phosphorylation levels detected in the salt sensitive *fer-4* and *herk1the1-4* mutants, corresponds with higher salt induced gene expression and a higher seedling sensitivity, suggesting that FER alone or the HERK1/THE1-4 combination is required for negatively regulating this branch of the salt induced signaling pathway. The shared oversensitive salt stress responses might suggest that the HERK1/THE1-4 combination could be required to regulate FER function. While calcium application alleviates early salt responses in an MPK6-dependent manner, PME inhibitor treatment can suppress also the detected mild halotropic root bending response observed in the over sensitive *fer-4* and *herk1the1-4* mutants. It remains to be understood why the effects of calcium or EGCG application on regulating salt induced pectin de-methyl esterification seem to be very similar. Further investigations will be required to understand the molecular mode of action and the role of CrRLK1Ls in detecting cell wall changes induced upon salt stress, which we show here to be responsible for a major part of salt stress responses in Arabidopsis.

## Materials and methods

### Plant materials and growth condition

*Arabidopsis thaliana* ecotype Col-0 was used as wildtype. Experiments were conducted on several mutants listed in Table S1. Before sowing, seeds were surface sterilized with 50% bleach for 10 min and washed 3 times with sterile distilled MilliQ water. The seeds were sown after two days of vernalization at 4°C in the dark. For MPK6 phosphorylation assays and gene expression analysis 15 seeds were germinated and grown in 2 mL half-strength MS medium supplemented with 0.1% 2(N-morpholino) ethane sulphonic acid (MES) buffer and 0.5% sucrose (pH 5.7) in 12 well multi-well plates. Chemicals used for the analysis of calcium effects on salt induced responses were in distilled sterile MilliQ water (used as mock), 5 mM CaCl_2_ (in sterile water), 100 mM NaCl (in sterile water) or CaCl_2_/NaCl combinations. Similarly, chemicals used for the analysis of PME inhibition were DMSO (mock), 50 μM EGCG (Sigma-Aldrich; in DMSO), 100 mM NaCl or a combination of EGCG/NaCl. All these experiments were performed on 7-day old seedlings. All treatments contained equal amounts of DMSO. For the halotropism assays, the media used were made with the recipe stated above and supplemented with 1% agar. Seedlings were grown in 70-degree holders and after 4 days the lower part of the media under a diagonal line was cut with sterile thin glass slide and replaced with 1/2 MS supplemented with H_2_O (mock) or 200 mM NaCl. In the case of halotropism assays performed in the presence of EGCG the agar strength of the medium was enhanced to 1.25% to avoid root tip growth into the medium. In these experiments lower gradients contained DMSO (mock), 50 μM EGCG, 200 mM NaCl or a combination of EGCG/NaCl. All treatments contained equal amounts of DMSO. All results are based on independent experiments and number of biological replicates are stated in figure legends or are shown in supplemental figures. An Epson Scanner was used to image plates at 24 h or 48 h after treatment. For cell wall extraction analysis and ion content measurements seedlings were grown for 6 days in 250 mL Erlenmeyer flasks containing 110 mL of 1/2 MS medium, 0.1% MES and 1% sucrose (pH 5.7) on a flask shaker at the constant speed of 130 rotations per minute. Treatments were performed with H_2_O, or 100 mM NaCl and samplings done after 24 h. For time-lapse growth analysis seedlings were grown in 1/2 MS medium, 0.1% MES (pH 5.7) supplemented with 1% agar in square plates positioned in vertical holders (90 degrees). After 7 days, seedlings were transferred in mock (H_2_O) or 125 mM NaCl containing plates and pictures taken, with an automatized system every 20 minutes for over 42 h. For the beaching assays seeds were germinated and grown for 10 days in horizontally oriented square plates containing 1/2 MS medium, 0.1% MES, 1% agar (+/− 150 mM NaCl) pH 5.7. Seedlings were grown in long-day conditions (16 h light, 22°C/8 h dark, 18°C) at 150 μmol m^-2^ s^-1^ photon flux density.

### Salt stress assays

Analysis of root halotropic responses was performed after scanning seedlings grown in square plates with an Epson scanner. Roots were traced with SmartRoot 2.0, an open-source plugin for Fiji/ImageJ (https://fiji.sc) and root directions were used to determine root bending (angle). For time-lapse based root assays, an automated script based on root edge detection was developed (https://github.com/jasperlamers/timelapse-backtracing) and used to detect and trace root length using the last picture taken. For salt tolerance assays, plants were grown in pots under short light photoperiod (8 h light/16 h dark) in growth chambers set at 21 ºC and with controlled humidity (70%). Ten days after germination (1-week old seedlings) plants were transferred to new pots saturated with 0 or 75 mM of NaCl solution. For 3 weeks plants were watered twice a week with MilliQ water. Rosette fresh weights were measured at the end of the 3 weeks. The experiments were repeated 2 times with similar results.

### Gene expression analysis and MPK6 phosphorylation assays

7-day old seedlings grown as stated above, were treated with mock or 100 mM NaCl. After 15 minutes (MPK6 assays) or 1 h (gene expression analysis), seedlings were flash frozen in 2 mL tubes containing metal beads in liquid nitrogen and ground into a fine powder by using a paint shaker. Total RNA was isolated using EasyPrep Plant Total RNA Extraction Miniprep (Bioland). DNAse treatment were performed as in (Gigli-Bisceglia et al., 2018; Savatin et al., 2014) and cDNA synthetized with iScript cDNA Synthesis Kit (Biorad) according to the manufacturer’s instructions. qRT-PCR analysis was conducted by CFX96 Real-Time System (Biorad) using qPCRBIO SyGreen Blue Mix Lo-ROX (Sopachem) master mix and primers (Table S2) diluted according. Gene expression analyses were performed as in (Engelsdorf et al., 2018; Gigli-Bisceglia et al., 2018) and are shown as relative expression to *ACT2*, used as reference gene. For protein extraction, the following extraction buffer was used; 50 mM Tris-HCl pH 7.5, 200 mM NaCl, 1 mM EDTA, 10% (v/v) glycerol, 0.1% (v/v) Tween 20, 1 mM PMSF, 1 mM DTT, 1× protease inhibitor cocktail (Sigma, P9599), 1x phosphatase inhibitor (Sigma, P2850). After 15 min incubation in ice, samples were centrifuged at 12000 rpm at 4°C for 15 min. The supernatants were used for western blot analysis. Protein concentration was assayed by Bradford assay (Biorad). 15 μg of total proteins were loaded on 10% SDS-PAGE gel. The samples were mixed with 4× loading buffer (240 mM Tris pH 6.8, 8% (w/v) SDS, 40% glycerol, 5% β-mercaptoethanol, 0.04% (w/v) bromophenol blue) and heated for 2 min at 95°C. Protein samples were separated and then transferred to the nitrocellulose membrane in Trans-Blot Turbo Transfer System (Biorad) at 25 V, 1.3 A for 7 min. After blocking with Tris Buffered Saline with Tween 20 (TBST) containing 5% (w/v) bovine serum albumin (BSA) (Sigma, A3608) for 1 h, the membrane was incubated overnight with the primary phospho-p44/42 MAPK (Erk1/2) (Thr202/Tyr204) antibody (Cell Signaling Technology, #9101). Subsequently, the membrane was washed 3 times with TBST and then incubated with secondary antibody, anti-rabbit IgG, HRP-linked antibody (Cell Signaling Technology, #7074) for 1 h. After washing 3 times with TBST the chemiluminescent was conducted by adding Clarity Western ECL Substrate (Biorad, #1705061) for 2 min and detected by ChemiDoc imaging system (Biorad). For equal loading analysis, membranes were stripped by stripping solution (100 mM Tris at pH 6.8, 2% SDS, 100 mM β-mercaptoethanol) for 1 h, and blocked as described previously incubation with β-Actin (C4) antibody (Santa Cruz biotechnology, sc-47778) for 1 h. Detection of chemiluminescence was performed as mentioned before.

### Cell wall extraction and dot blot assays

After 24 h treatment (see above for details) seedlings were snap frozen in liquid nitrogen and ground into a fine powder with a paint shaker. At least 3 biological replicates for each treatment were used. Cell wall material (AIR) was extracted as in (Gigli Bisceglia et al., 2018). After an overnight of drying, AIRs were weighed and sterile distilled MilliQ water (ph 7.2), used to extract pectins. Approximately 2-to 3 mg of AIR were boiled with 250 μl water for 1 h. After a short centrifuge (15 min, 13.000xg), supernatants were collected, and samples boiled again for an additional hour with 250 μl water. After 1h supernatants deriving from the same samples were pooled. Total sugar content was measured with the phenol/H_2_SO_4_ based method (Dubois) using galacturonic acid as standard (Dubois et al., 1951). Prior spotting on nitrocellulose membrane, 5 μg total sugars were adjusted to 5 μl by adding (if necessary) distilled MilliQ water (ph 7.2). For positive and negative control, the same amount of sugars were incubated for 1 h with NaCO3 pH 11 to remove pectin methyl esterification. Membranes were dried overnight at room temperature. Blocking (5% milk in PBS) was performed for 1 h. Next, membranes were incubated for 1 h with primary antibodies [LM20, 19 and 2F4, (http://www.plantprobes.net/index.php)] in the following buffers/concentrations; 1:250 in 5% milk powder in 1 × PBS for both Lm20/19 and 1:250 in 5% milk in T/Ca/S buffer (Tris-HCl 20 mM pH 8.2, CaCl2 0.5 mM, NaCl 150 mM) (http://www.plantprobes.net/index.php) (Verhertbruggen et al., 2009). After PBST washes (3 times) membranes were incubated with HRP-conjugated secondary antibodies (rat (IgM)/LM20 and LM19 mouse (IgG)/2F4 (in 5% milk PBS) and after washing chemiluminescence induced with Clarity Western ECL Substrate (Biorad, #1705061) and detected with ChemiDoc imaging system (Biorad).

### Statistical analysis

IBM SPSS Statistics v24 was used to perform statistical significance (Student’s t-test, ANOVA). Statistically significant differences are indicated with different letters for the one-way ANOVA/Tukey’s HSD test at α=0.05. Pairwise comparisons, and statistically significant differences according to Student’s T-test are represented with * P<0.05, ** P<0.01, *** P<0.001. For root time-lapse growth analysis, root elongation rate was calculated over 2 h intervals, averaged per hour and expressed either as a ratio between treatment vs controls (Figure 2A) or as growth rate (cm/ h) (Figure S3A). Each curve comprises several dots, representing the average growth rate and the corresponding standard error analyzed in each of the time lapse images containing 5 seedlings per plates (only correctly segmented roots were included, n>12). In this assay statistical comparisons were performed by analyzing every timepoints for equal variance using a Levene’s test, followed by T-test to compare different genotypes at the same timepoint. The significance level (p= 0.05) was corrected for multiple testing by using stepwise Sidak correction.

## Acknowledgments

We thank Paul Knox for providing the probes used in this paper and Hermann Höfte for sharing with us *the1-1 fer-4* line. This work was supported by the European Research Council (ERC) under the European Union’s Horizon 2020 research and innovation programme (grant agreement No 724321; ERC CoG grant Sense2SurviveSalt awarded to C.T.).

## Author Contributions

N.G.B conceived the presented idea and performed most of the experiments of the study. C.T and N.G.B have designed the work. E.VZ provided the time-lapse based root growth measurements and the statistical analysis associated to that. J. L. developed the automated software to analyze images from the time-lapse setup and assisted with halotropism assays. W.H performed the halotropism assays with salt and EGCG combinations. C.T supervised the findings and discussed the results with N.G.B., together they wrote the manuscript, with help of all authors.

## Supplemental Figure Legends

**Figure S1. A.** Seeds of Col-0, *the1-1, the1-4, herk1, herk1the1-4, fer-4* and *the1-1fer-4* were germinated and grown for 10 days in 0.5% MS agar plates containing 150 mM NaCl. Similar results have been obtained in 4 different biological replicates. 7-day old seedlings grown in liquid 0.5% MS medium supplemented with 0.5% sucrose were treated for 15 minutes with H_2_0 (mock, m) or 100 mM NaCl (N). MPK6 phosphorylation was detected in Col-0, *mpk6-3* **(B)** and in Col-0, *herk1, the1-4, herk1the1-4* and *fer-4* (**C**) by immunoblot using specific antibody against the phosphorylated forms of both MPK3 and MPK6 (*α-*p44/42). Equal loading is indicated with Ponceau Red staining or upon stripping on the same membrane by using *β*-actin Arabidopsis specific antibody. **D.** Seedlings grown as in B/C were treated for 1 h with H_2_0 (mock) or 100 mM NaCl (NaCl) to analyze the expression levels of selected salt-induced marker genes. *ProPEP3, WRKY40, RRTF1* expression relative to *ACT2* was determined by qRT-PCR. Values represent means with error bars indicating SD. One-way ANOVA and Tukey’s HSD (*α*= 0.05) was performed on different genotypes within the same treatment (black letters for mock comparisons, red for NaCl treatments). n= 3.

**Figure S2. A.** The 3 biological replicates pooled in figure 2B are shown (24 h, upper panel, 48 h lower panel). Bars indicating standard errors (SE). Letters indicate statistically significant differences according to one-way ANOVA and Tukey’s HSD (*α*= 0.05) between genotypes within one treatment condition (n= 50) (black letters= mock, red= NaCl). **B.** Root elongation analyzed 24 h and 48 h after mock or NaCl gradient application in experiment presented in figure 2B. Bars indicate standard deviation. Letters indicate statistically significant differences according to one-way ANOVA in 3 independent experiments and Tukey’s HSD (*α*= 0.05) between genotypes within one treatment condition (black letters= mock, red= NaCl). **C.** 4-day old seedlings of Col-0, *mpk6-3* were treated as in figure 2B and root angle was analyzed after 24 h and 48 h. Histogram represents the average root angle of 2 independent experiments, while bars show the standard errors (SE). (n= 45). Asterisks show statistical comparisons between the same treatment in different genotypes; ***P < 0.001, *n.s.* not significant. **D.** Representative rosette size of 4-week-old Col-0, *fer-4, herk1the1-4* plants grown in short day conditions and watered once with either MilliQ water (mock) or 75 mM NaCl (NaCl). Plants were photographed after 3 weeks of MilliQ watering. Rosette fresh weight reported in mg (**E**) and salt response (**F**) are shown. In E, values show means with error bars indicating standard errors (SE). Letters indicate statistically significant differences according to one-way ANOVA and Tukey’s HSD (*α*= 0.05) between genotypes within one treatment condition (lowercase letters= mock, capitalized letters= NaCl). The experiment has been repeated twice with similar results. In F, bars represent a ratio between NaCl treated genotypes and the corresponding genotype in control conditions. Error bars are SE. *n.s.* = not significant based on Student’s T-test (P, <0.05).

**Figure S3. A.** Root elongation rates (expressed in cm/h) of the two independent biological replicates pooled in figure 3A are calculated as the average elongation over 2 h intervals in a 42 h time course experiment. Dots represent average elongation lengths and corresponding standard errors (n= 15 per treatment, per replicate). **B.** Biological replicates of the experiment shown in figure 3B. **C.** 4-day old seedlings of Col-0, *fer-4* and *herk1the1-4* were subjected to mock, 200 mM NaCl, 10 mM CaCl_2_ or NaCl/CaCl_2_ gradients. Root angle (direction) at 24 h and 48 h is shown. Histogram represents the average root angle of 2 independent experiments, while bars show the average of standard errors (SE). One-way ANOVA and Tukey’s HSD (*α*= 0.05) was used to analyze statistical differences between treatments within the same genotypes. Asterisks show statistical comparisons between the same treatment in different genotypes; ***, P < 0.001, *, P< 0.05 (n=45). **D.** Root elongation analyzed after 24 h and 48 h experiment shown in C. Histogram represents the average root length of 2 independent experiments, while bars show standard deviation (SD). Statistical analysis has been performed as in C. (n= 45).

**Figure S4. A.** Schematic representation of the cell wall epitopes recognized by the monoclonal antibodies used in this study. **B.** 5 *μ*g total sugars derived from Col-0, *fer-4* and *herk1the1-4* mutants or experiment presented in Figure 4B were spotted on nitrocellulose membranes after 1 h incubation with ddH_2_O (native) or Na_2_CO_3_ pH 11 (deM-HG). Dotblot was performed with LM20. Three biological replicates are shown. **C.** Biological replicate for figure 4C is shown. **D/E.** Biological replicates of Figure 4E. Root angles analyzed at 24 h **(D)** or at 48 h (**E)** are plotted as average direction of 25 seedlings each treatment. Bars show standard error (SE). One-way ANOVA and Tukey’s HSD (*α*= 0.05) was used to analyze statistical differences between treatments within the same genotypes. **F**. Root elongation of experiments shown in Figure 4E, analyzed after 24 h (left) and 48 h (right) in 4-day old seedlings of Col-0, *fer-4* and *herk1the1-4* treated with with DMSO (mock), 100 mM NaCl (NaCl), 50 *μ*M EGCG (EGCG) and NaCl/EGCG (NaCl/EGCG) gradients. Histogram represents the average root length of 3 independent experiments shown in D and E, while bars show standard deviation (SD). Statistical analysis has been performed as in C. (n= 75). **G.** 5 *μ*g total sugars derived from Col-0 experiments presented in Figure 4F (left) or 4G (right) were spotted on nitrocellulose membranes after 1h incubation with ddH_2_O (native) or Na_2_CO_3_ pH 11 (deM-HG). Dot-blot was performed with LM20. Three biological replicates are shown.

