## Supplemental Tables and Figures for "Arabidopsis root responses to salinity depend on pectin modification and cell wall sensing"

**Table S1. Arabidopsis genotypes used in this study.**

| Genotype | AGI | Allele | Background | Reference |
|---|---|---|---|---|
| *the1-1* | *AT5G54380* | *the1-1* | Col-0 | (Hématy et al., 2007) |
| *the1-4* | *AT5G54380* | *the1-4* | Col-0 | (Li et al., 2009) |
| *herk1* | *AT3G46290* | *herk1* | Col-0 | (Li et al., 2009) |
| *fer-4* | *AT3G51550* | *fer-4* | Col-0 | (Duan et al., 2010) |
| *the1-1 fer1-4* | *AT5G54380, AT3G51550* | *the1-1 fer1-4* | Col-0 | (Gonneau et al., 2018) |
| *mpk6-3* | *AT2G43790* | *mpk6-3* | Col-0 | (Liu and Zhang, 2004) |

**Table S2. Primers used in this study.**

| Name | AGI | sequence (5'-3') | Reference |
|---|---|---|---|
| *ACT2_for* | AT3G18780 | CTTGCACCAAGCAGCATGAA | (Czechowski et al., 2005) |
| *ACT2_rev* | | CCGATCCAGACACTGTACTTCCTT | |
| *PROPEP3_for* | AT5G64905 | CAACGATGGAGAATCTCAGA | (Engelsdorf et al., 2018; Gigli-Bisceglia et al., 2018) |
| *PROPEP3_rev* | | CTAATTGTGTTTGCCTCCTTT | |
| *WRKY40 _for* | AT1G80840 | GATCCACCGACAAGTGCTTT | (Denoux et al., 2008) |
| *WRKY40_rev* | | AGGGCTGATTTGATCCCTCT | |
| *RRTF1-for* | AT4G34410 | TATAGGAGCAAAGGCAAGTGCA | |
| *RRTF1_rev* | | ACTCCTCCATATTGCAATCCCC | |

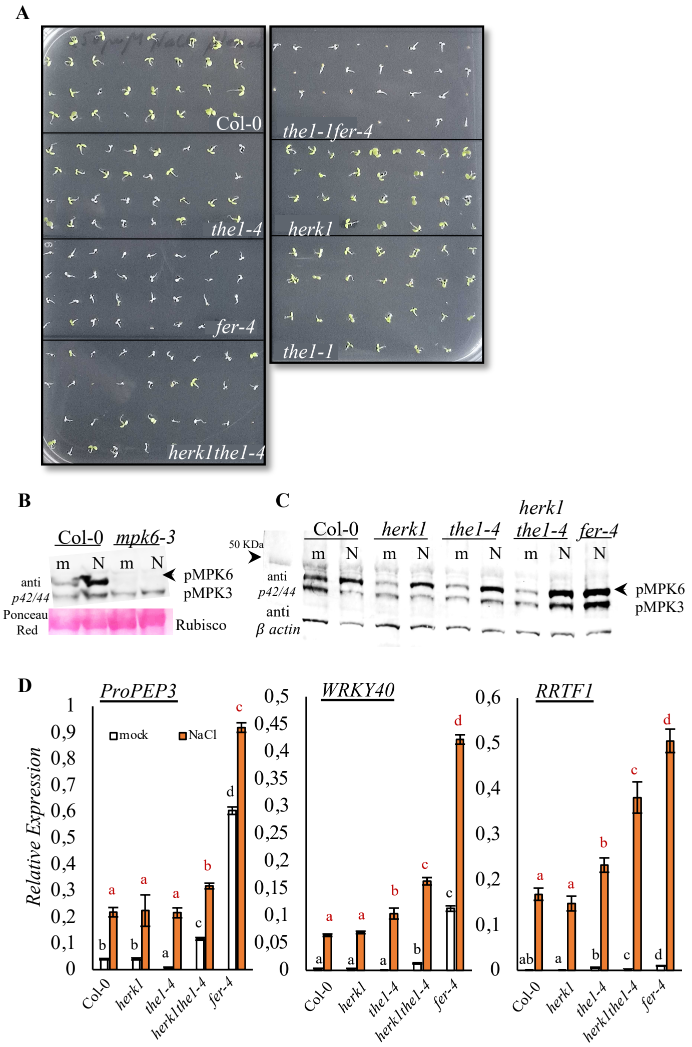


**Figure S1. A.** Seeds of Col-0, *the1-1, the1-4, herk1, herk1the1-4, fer-4* and *the1-1fer-4* were germinated and grown for 10 days in 0.5% MS agar plates containing 150 mM NaCl. Similar results have been obtained in 4 different biological replicates. 7-day old seedlings grown in liquid 0.5% MS medium supplemented with 0.5% sucrose were treated for 15 minutes with $H_20$ (mock, m) or 100 mM NaCl (N). MPK6 phosphorylation was detected in Col-0, *mpk6-3* **(B)** and in Col-0, *herk1, the1-4, herk1the1-4* and *fer-4* (**C**) by immunoblot using specific antibody against the phosphorylated forms of both MPK3 and MPK6 ($\alpha$-p44/42). Equal loading is indicated with Ponceau Red staining or upon stripping on the same membrane by using $\beta$-actin Arabidopsis specific antibody. **D.** Seedlings grown as in B/C were treated for 1 h with $H_20$ (mock) or 100 mM NaCl (NaCl) to analyze the expression levels of selected salt-induced marker genes. *ProPEP3, WRKY40, RRTF1* expression relative to *ACT2* was determined by qRT-PCR. Values represent means with error bars indicating SD. One-way ANOVA and Tukey's HSD ($\alpha$= 0.05) was performed on different genotypes within the same treatment (black letters for mock comparisons, red for NaCl treatments). n= 3.

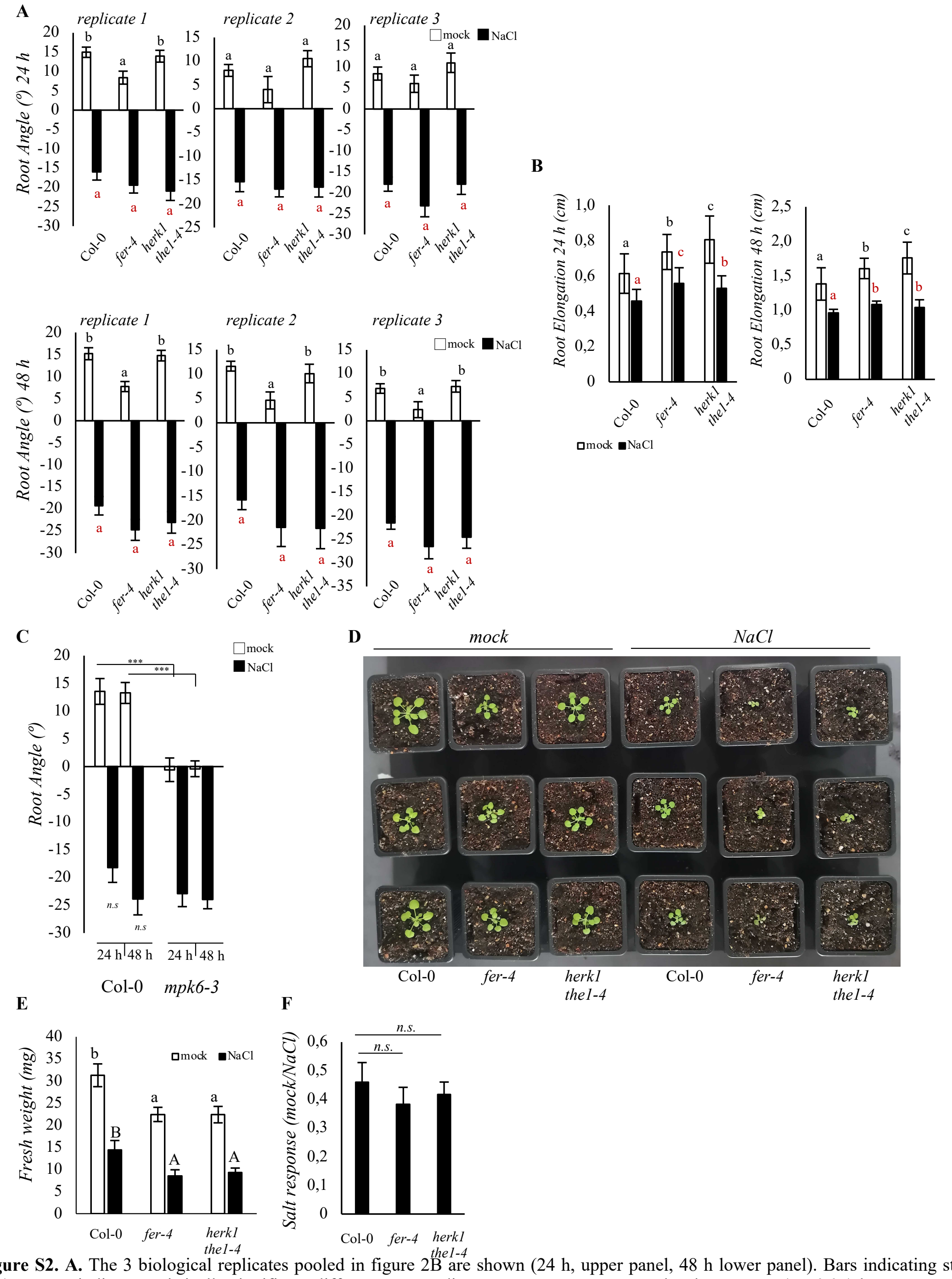


**Figure S2. A.** The 3 biological replicates pooled in figure 2B are shown (24 h, upper panel, 48 h lower panel). Bars indicating standard errors (SE). Letters indicate statistically significant differences according to one-way ANOVA and Tukey's HSD ($\alpha$= 0.05) between genotypes within one treatment condition (n= 50) (black letters= mock, red= NaCl). **B.** Root elongation analyzed 24 h and 48 h after mock or NaCl gradient application in experiment presented in figure 2B. Bars indicate standard deviation. Letters indicate statistically significant differences according to one-way ANOVA in 3 independent experiments and Tukey's HSD ($\alpha$= 0.05) between genotypes within one treatment condition (black letters= mock, red= NaCl). **C.** 4-day old seedlings of Col-0, *mpk6-3* were treated as in figure 2B and root angle was analyzed after 24 h and 48 h. Histogram represents the average root angle of 2 independent experiments, while bars show the standard errors (SE). (n= 45). Asterisks show statistical comparisons between the same treatment in different genotypes; ***P < 0.001, *n.s.* not significant. **D.** Representative rosette size of 4-week-old Col-0, *fer-4, herk1the1-4* plants grown in short day conditions and watered once with either MilliQ water (mock) or 75 mM NaCl (NaCl). Plants were photographed after 3 weeks of MilliQ watering. Rosette fresh weight reported in mg (**E**) and salt response (**F**) are shown. In E, values show means with error bars indicating standard errors (SE). Letters indicate statistically significant differences according to one-way ANOVA and Tukey's HSD ($\alpha$= 0.05) between genotypes within one treatment condition (lowercase letters= mock, capitalized letters= NaCl). The experiment has been repeated twice with similar results. In F, bars represent a ratio between NaCl treated genotypes and the corresponding genotype in control conditions. Error bars are SE. *n.s.* = not significant based on Student's T-test (P, <0.05).

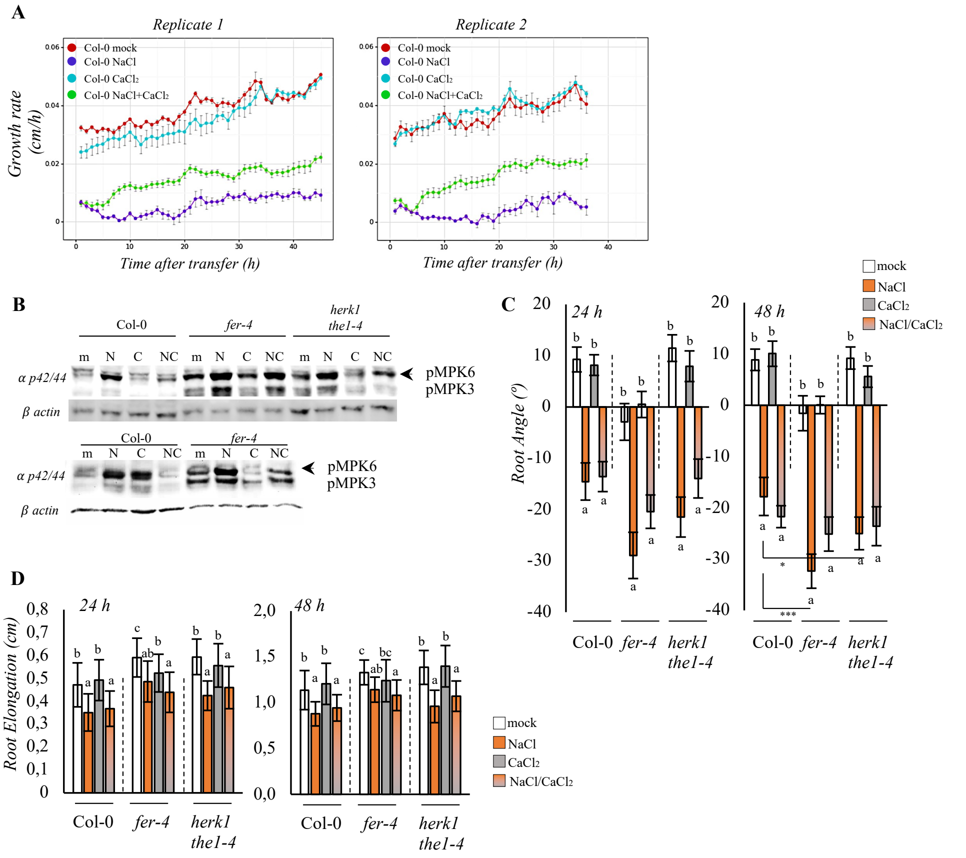


**Figure S3. A.** Root elongation rates (expressed in cm/h) of the two independent biological replicates pooled in figure 3A are calculated as the average elongation over 2 h intervals in a 42 h time course experiment. Dots represent average elongation lengths and corresponding standard errors (n= 15 per treatment, per replicate). **B.** Biological replicates of the experiment shown in figure 3B. **C.** 4-day old seedlings of Col-0, *fer-4* and *herk1the1-4* were subjected to mock, 200 mM NaCl, 10 mM $CaCl_2$ or NaCl/$CaCl_2$ gradients. Root angle (direction) at 24 h and 48 h is shown. Histogram represents the average root angle of 2 independent experiments, while bars show the average of standard errors (SE). One-way ANOVA and Tukey's HSD ($\alpha$= 0.05) was used to analyze statistical differences between treatments within the same genotypes. Asterisks show statistical comparisons between the same treatment in different genotypes; ***, P < 0.001, *, P< 0.05 (n=45). **D.** Root elongation analyzed after 24 h and 48 h experiment shown in C. Histogram represents the average root length of 2 independent experiments, while bars show standard deviation (SD). Statistical analysis has been performed as in C. (n= 45).

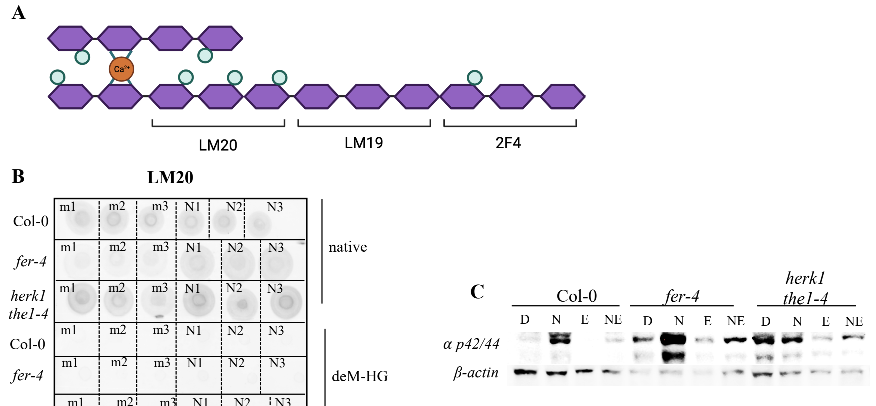


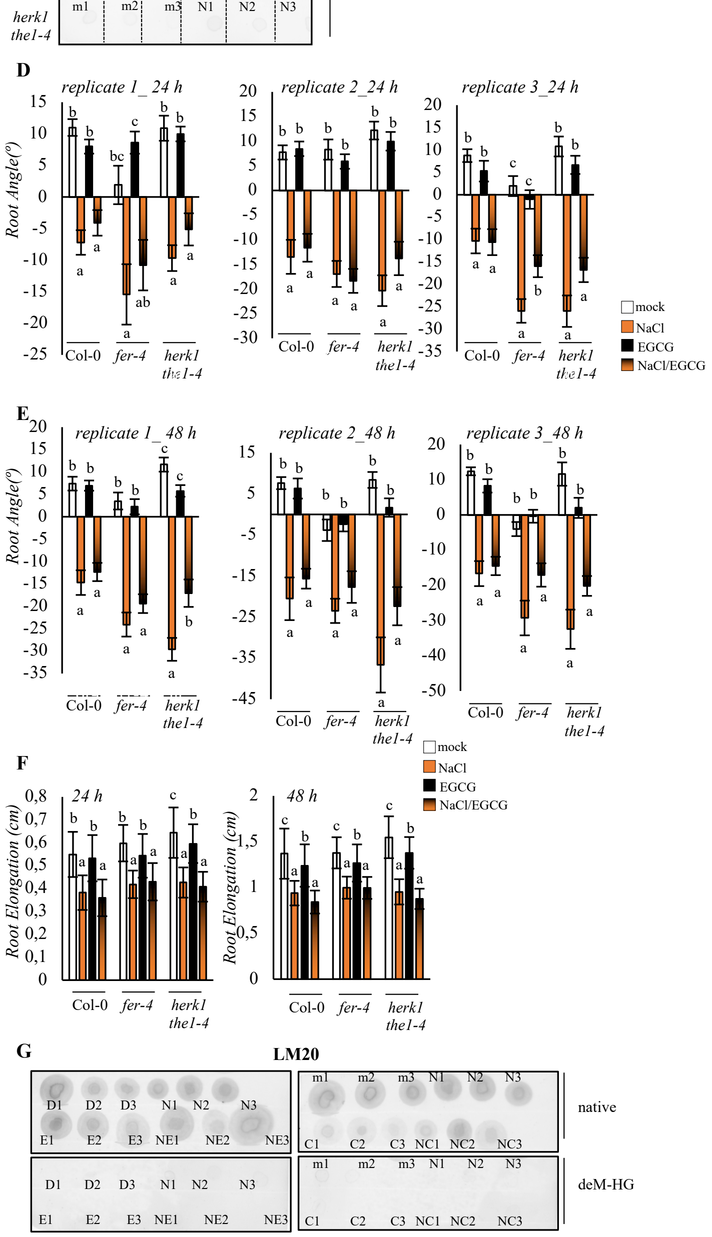


**Figure S4. A.** Schematic representation of the cell wall epitopes recognized by the monoclonal antibodies used in this study. **B.** 5 $\mu$g total sugars derived from Col-0, *fer-4* and *herk1the1-4* mutants or experiment presented in Figure 4B were spotted on nitrocellulose membranes after 1 h incubation with ddH$_2$O (native) or Na$_2$CO$_3$ pH 11 (deM-HG). Dot-blot was performed with LM20. Three biological replicates are shown. **C.** Biological replicate for figure 4C is shown. **D/E.** Biological replicates of Figure 4E. Root angles analyzed at 24 h **(D)** or at 48 h **(E)** are plotted as average direction of 25 seedlings each treatment. Bars show standard error (SE). One-way ANOVA and Tukey's HSD ($\alpha$= 0.05) was used to analyze statistical differences between treatments within the same genotypes. **F.** Root elongation of experiments shown in Figure 4E, analyzed after 24 h (left) and 48 h (right) in 4-day old seedlings of Col-0, *fer-4* and *herk1the1-4* treated with with DMSO (mock), 100 mM NaCl (NaCl), 50 $\mu$M EGCG (EGCG) and NaCl/EGCG (NaCl/EGCG) gradients. Histogram represents the average root length of 3 independent experiments shown in D and E, while bars show standard deviation (SD). Statistical analysis has been performed as in C. (n= 75). **G.** 5 $\mu$g total sugars derived from Col-0 experiments presented in Figure 4F (left) or 4G (right) were spotted on nitrocellulose membranes after 1h incubation with ddH$_2$O (native) or Na$_2$CO$_3$ pH 11 (deM-HG). Dot-blot was performed with LM20. Three biological replicates are shown.
